# The mechanical and sensory signature of plant-based and animal meat

**DOI:** 10.1101/2024.04.25.591207

**Authors:** Skyler R. St. Pierre, Ethan C. Darwin, Divya Adil, Magaly C. Aviles, Archer Date, Reese A. Dunne, Yanav Lall, María Parra Vallecillo, Valerie A. Perez Medina, Kevin Linka, Marc E. Levenston, Ellen Kuhl

## Abstract

Eating less meat is associated with a healthier body and planet. Yet, we remain reluctant to switch to a plant-based diet, largely due to the sensory experience of plant-based meat. Food scientists characterize meat using a double compression test, which only probes one-dimensional behavior. Here we use tension, compression, and shear tests–combined with constitutive neural networks–to automatically discover the behavior of eight plant-based and animal meats across the entire three-dimensional spectrum. We find that plant-based sausage and hotdog, with stiffnesses of 95.9±14.1kPa and 38.7±3.0kPa, successfully mimic their animal counterparts, with 63.5±45.7kPa and 44.3±13.2 kPa, while tofurky is twice as stiff, and tofu is twice as soft. Strikingly, a complementary food tasting survey produces in nearly identical stiffness rankings for all eight products (***ρ*** =0.833, p=0.015). Probing the fully three-dimensional signature of meats is critical to understand subtle differences in texture that may result in a different perception of taste.

Our data and code are freely available at https://github.com/LivingMatterLab/CANN

---

Current meat production is inefficient and unsustainable [1]. It is a key driver for climate change, environmental degradation, and antibiotic resistance [2]. A common strategy to quantify the efficiency of animal meat is to compare energy out versus energy in, and protein out versus protein in [3]. On a global average, energy conversion efficiencies range from 11% for poultry to 10% for pork, and 1% for beef and sheep; while protein conversion efficiencies range from 20% for poultry to 15% for pork, 4% for beef, and 3% for sheep [4]. In other words, cattle are a particularly inefficient conversion system: They convert only one percent of the gross energy and four percent of the protein in their feed to energy and protein in edible beef [5].

At the same time, the consumption of a kilogram of beef is estimated to have the same greenhouse gas emission impact as driving 1.172km in a gas-fueled passenger car [6]. On a global average, the estimated greenhouse gas emission in kg carbon dioxide equivalents per kg food varies from 100 for beef to 40 for sheep, 12 for pork, and 10 for poultry, while it is only 3 for tofu [7]. These alarming numbers underscore that the global food system is a major source of greenhouse gas emissions [8]. In fact, experts estimate that it will be critical to change the global food system to meet the Paris Agreement of limiting the increase in global temperature to 1.5 to 2.0°C above its pre-industrial levels [9].

Equally concerning is the increasing use of antimicrobials in food animals to promote growth, increase health, and maintain food safety [10]. On a global average, the estimated annual consumption of antimicrobials per kilogram of animal meat ranges from 172mg for pigs, to 148mg for chicken, and 45mg for cattle [11]. These practices contribute to the spread of drug-resistant pathogens in both livestock and humans. Based on current trends, medical experts expect 10 million annual deaths from antimicrobial resistance in 2050, a 14-fold increase over current deaths [12]. Beyond these global concerns, eating red and processed meat is associated with an increased individual risks of colorectal cancer, chronic kidney disease, type-2 diabetes, and cardiovascular disease [13]. Taken together, it is becoming increasingly clear that a shift towards consuming less meat is needed for the health of the planet and people [1]

Plant-based meat products are alternatives to animal meat that are produced directly from plants [14]. The fundamental idea is to avoid using animals to convert vegetation into meat. Recent studies suggest that plant-based meat products have a 50% lower environmental impact than animal meat [15]; yet, their adoption into our diets remains low [16]. Out of all global greenhouse emissions related to the production of food, animal food contributes 57%, while plant-based food contributes only 29% [17]. In fact, the livestock industry alone is estimated to contribute 12-18% to the total greenhouse gas emissions; it decreases water quality, and increases water scarcity [13]. Although more and more people are intrigued by a flexitarian diet and intentionally reduce their animal meat intake, most of us do not eliminate meat entirely [18]. A recent consumer survey revealed that in the United States, only one third of participants were very or extremely likely to buy plant-based meat products [19]. This reluctancy supports the emerging view that the key to a more sustainable food system is not to convince consumers to give up the foods they like [1]; instead, scientists should focus on creating plant-based products that taste the same or better than animal meat [20]. In fact, our sensory experience–including taste and texture–is one of the most important reasons for choosing which product we eat [16]. But how can we reliably and reproducibly quantify and compare the texture of different meat products?

According to its widely-accepted definition, texture is the sensory and functional manifestation of the structural, mechanical, and surface properties of food. [21]. *Sensory manifestation* has been the focus of numerous food texture surveys [22], but their results vary greatly by personal preference, subjective interpretation, cultural background, sensory ability, lack of standardization, and various external factors [16]. *Functional manifestation* has traditionally been quantified by texture profile analysis [23], using a double mechanical compression test [24]. This method is quantitative, objective, and semi-standardized [25], but only provides insights into the one-dimensional compressive behavior of the product [26]. If plant-based meat products are to mimic the true textural properties of animal meat, how can we comprehensively characterize the fully three-dimensional mechanics of plant-based and animal meat? And, ultimately, how can we use this knowledge to inform the design of more animal-like meat alternatives?

## The objective of this study is to understand the mechanical signatures of plant-based and animal meat

Animal meat consists of muscle, connective tissue, fat, and water, and is dominated by its anisotropic microstructure [27]. In contrast, most plant-based meats are made of soy-, wheat-, or pea-based proteins, they have no pronounced fibers, and are generally isotropic [14]. Here, to unify the comparison, we focus on processed meat products– plant-based and animal sausage and hotdog–and assume that these products no longer have fibers, but are all *homogeneous* and *isotropic*. For comparison, we also characterize processed plant-based and animal turkey, and two plain soy-protein products, extrafirm and firm tofu. We systematically compare all eight products through standardized mechanical tension, compression, and shear tests [28], combined with automated model discovery, to discover the best models and parameters for each product [29], and conduct a food texture survey [21] with *n* = 16 participants to explore whether the results align with our sensory perception. To accelerate discovery and innovation in plant-based food technologies [1], we share all raw data, methods, algorithms, and results on a public open source discovery platform.

## Results

### Mechanical testing

We mechanically test eight products, five plant-based, tofurky, plant-based sausage, plant-based hotdog, extrafirm tofu, and firm tofu, and three animal-based, spam turkey, animal sausage, and animal hotdog. Table 1 summarizes the brand name, manufacturer, and list of ingredients of all eight products. For each product, we perform tension, compression, and shear tests on at least *n* = 5 samples per testing mode. Table 2 summarizes the means ± standard error of the means for all eight products and all three tests.

**Table 1.** Plant-based and animal meat products. Products; brands; ingredients; stiffnesses in tension, compression, shear, and mean stiffness; peak strain and stress; discovered best-in-class one- and two-term models, parameters, and goodness of fit; and texture features soft, hard, chewy, moist, fatty, and meat-like.

|  | TF tofurky | PS plant sausage | ST spam turkey | AS animal sausage |
| --- | --- | --- | --- | --- |
| **brand** | Ham-Style Roast Tofurky<br>Hood River, OR | Vegan Frankfurter Sausage<br>Field Roast<br>Seattle, WA | Spam Oven Roasted Turkey<br>Spam, Hormel Foods Co<br>Austin, MN | Turkey Polska Kielbasa<br>Hallshire Farm<br>New London, WI |
| **ingredients** | water, vital wheat gluten, tofu (water, soybeans, magnesium chloride, calcium chloride), expeller pressed canola oil; contains 2% or less: sea salt, spices, granulated garlic, cane sugar, natural flavors, natural smoke flavor, color, oat fiber | filtered water, vital wheat gluten, expeller pressed safflower oil, organic expeller pressed palm fruit oil, barley malt, naturally flavored yeast extract, tomato paste, apple cider vinegar, paprika, sea salt, onions, spices, whole wheat flour | white turkey, turkey broth, salt, modified potato starch, sugar, dextrose, sodium nitrite | turkey, mechanically separated turkey, water, corn syrup; contains 2% or less: natural flavors, salt, dextrose, oat fiber, modified corn starch, monosodium glutamate, sodium erythorbate, yeast extract, sodium nitrite |
| **stiffness** | $E_{ten} = 167.9 \pm 34.8$ kPa<br>$E_{com} = 222.8 \pm 33.6$ kPa<br>$E_{shr} = 224.5 \pm 38.6$ kPa<br>$E_{mean} = 205.1 \pm 32.2$ kPa | $E_{ten} = 79.9 \pm 33.0$ kPa<br>$E_{com} = 106.3 \pm 15.2$ kPa<br>$E_{shr} = 101.5 \pm 20.5$ kPa<br>$E_{mean} = 95.9 \pm 14.1$ kPa | $E_{ten} = 100.3 \pm 4.9$ kPa<br>$E_{com} = 72.2 \pm 8.9$ kPa<br>$E_{shr} = 69.9 \pm 7.9$ kPa<br>$E_{mean} = 80.8 \pm 16.9$ kPa | $E_{ten} = 115.5 \pm 13.2$ kPa<br>$E_{com} = 44.9 \pm 4.5$ kPa<br>$E_{shr} = 30.0 \pm 1.7$ kPa<br>$E_{mean} = 63.5 \pm 45.7$ kPa |
| **peak** | $\epsilon = 10\%$, $\sigma = 15.4$ kPa | $\epsilon = 15\%$, $\sigma = 10.5$ kPa | $\epsilon = 15\%$, $\sigma = 13.8$ kPa | $\epsilon = 15\%$, $\sigma = 17.6$ kPa |
| **one-term model** | $w_5 = 33.08$ kPa<br>$R^2 = 0.9810$ | $w_5 = 16.01$ kPa<br>$R^2 = 0.9588$ | $w_2 = 3.21$ kPa, $w_2^* = 3.92$<br>$R^2 = 0.8366$ | $w_2 = 0.42$ kPa, $w_2^* = 16.01$<br>$R^2 = 0.5310$ |
| **two-term model** | $w_1 = 12.27$ kPa<br>$w_5 = 20.94$ kPa<br>$R^2 = 0.9799$ | $w_1 = 6.56$ kPa<br>$w_5 = 9.34$ kPa<br>$R^2 = 0.9624$ | $w_1 = 11.33$ kPa<br>$w_3 = 51.77$ kPa<br>$R^2 = 0.8464$ | $w_1 = 3.63$ kPa<br>$w_4 = 11.85$ kPa, $w_4^* = 11.83$<br>$R^2 = 0.5311$ |
| **texture** | soft $1.7 \pm 0.9$ hard $4.0 \pm 0.8$<br>chewy $3.9 \pm 1.0$ moist $1.6 \pm 0.8$<br>fatty $1.7 \pm 0.8$ meat $2.5 \pm 0.9$ | soft $3.1 \pm 1.0$ hard $2.2 \pm 0.9$<br>chewy $3.7 \pm 0.9$ moist $2.3 \pm 1.0$<br>fatty $2.4 \pm 0.9$ meat $2.8 \pm 1.4$ | soft $4.3 \pm 0.6$ hard $1.3 \pm 0.5$<br>chewy $2.5 \pm 1.3$ moist $3.7 \pm 0.9$<br>fatty $3.6 \pm 1.1$ meat $3.7 \pm 1.4$ | soft $3.4 \pm 1.1$ hard $2.3 \pm 1.0$<br>chewy $3.3 \pm 1.0$ moist $4.1 \pm 0.9$<br>fatty $4.4 \pm 1.1$ meat $4.9 \pm 0.3$ |
|  | AH animal hotdog | PH plant hotdog | ET extrafirm tofu | FT firm tofu |
| **brand** | Classic Uncured Wiener<br>Oscar Mayer, Kraft Heinz Co<br>Chicago, IL | Signature Stadium Hot Dog<br>Field Roast<br>Seattle, WA | Organic Tofu extrafirm<br>House Foods<br>Garden Grove, CA | Organic Firm Tofu<br>365 by Whole Foods Market<br>Austin, TX |
| **ingredients** | mechanically separated chicken, mechanically separated turkey, pork, water, corn syrup; contains 2% or less: salt, sodium phosphate, potassium chloride, sodium diacetate, sodium benzoate, sodium ascorbate, flavor, sodium nitrate | water, soybean oil, pea protein, potato starch, contains 2% or less: methylcellulose, carrageenan, sea salt, brown rice protein, faba bean protein, garlic powder, beet powder, cane sugar, natural flavor, wheat gluten, soy protein, konjac flour, potassium chloride, xanthan gum, spices, paprika, red rice flour | water, organic soybeans, calcium sulfate, calcium chloride | water, organic soybeans, calcium sulfate, magnesium chloride |
| **stiffness** | $E_{ten} = 58.8 \pm 3.4$ kPa<br>$E_{com} = 41.0 \pm 5.2$ kPa<br>$E_{shr} = 33.0 \pm 4.5$ kPa<br>$E_{mean} = 44.3 \pm 13.2$ kPa | $E_{ten} = 39.6 \pm 4.3$ kPa<br>$E_{com} = 41.1 \pm 9.2$ kPa<br>$E_{shr} = 35.3 \pm 7.9$ kPa<br>$E_{mean} = 38.7 \pm 3.0$ kPa | $E_{ten} = 34.0 \pm 6.6$ kPa<br>$E_{com} = 24.4 \pm 9.5$ kPa<br>$E_{shr} = 24.1 \pm 5.7$ kPa<br>$E_{mean} = 27.5 \pm 5.6$ kPa | $E_{ten} = 26.5 \pm 3.7$ kPa<br>$E_{com} = 30.3 \pm 3.2$ kPa<br>$E_{shr} = 22.3 \pm 10.1$ kPa<br>$E_{mean} = 26.4 \pm 4.0$ kPa |
| **peak** | $\epsilon = 35\%$, $\sigma = 20.8$ kPa | $\epsilon = 20\%$, $\sigma = 7.5$ kPa | $\epsilon = 10\%$, $\sigma = 3.2$ kPa | $\epsilon = 15\%$, $\sigma = 3.5$ kPa |
| **one-term model** | $w_2 = 2.55$ kPa, $w_2^* = 2.59$<br>$R^2 = 0.9308$ | $w_2 = 2.61$ kPa, $w_2^* = 2.35$<br>$R^2 = 0.9743$ | $w_1 = 4.42$ kPa<br>$R^2 = 0.8318$ | $w_1 = 4.51$ kPa<br>$R^2 = 0.9494$ |
| **two-term model** | $w_1 = 5.80$ kPa<br>$w_7 = 22.83$ kPa<br>$R^2 = 0.9471$ | $w_1 = 6.22$ kPa<br>$w_3 = 7.25$ kPa<br>$R^2 = 0.9747$ | $w_1 = 4.42$ kPa<br>$R^2 = 0.8318$ | $w_1 = 4.51$ kPa<br>$R^2 = 0.9494$ |
| **texture** | soft $3.7 \pm 1.0$ hard $1.8 \pm 0.8$<br>chewy $3.3 \pm 0.9$ moist $3.8 \pm 1.0$<br>fatty $3.8 \pm 1.0$ meat $3.9 \pm 1.3$ | soft $3.4 \pm 0.9$ hard $2.2 \pm 0.9$<br>chewy $3.3 \pm 1.2$ moist $3.6 \pm 0.8$<br>fatty $3.0 \pm 1.4$ meat $3.2 \pm 1.3$ | soft $4.5 \pm 0.6$ hard $1.3 \pm 0.6$<br>chewy $2.9 \pm 1.3$ moist $4.1 \pm 1.0$<br>fatty $1.9 \pm 1.1$ meat $1.3 \pm 0.6$ | soft $4.8 \pm 0.4$ hard $1.0 \pm 0.3$<br>chewy $2.6 \pm 1.3$ moist $4.0 \pm 1.4$<br>fatty $1.5 \pm 0.8$ meat $1.2 \pm 0.4$ |

**Table 2.**
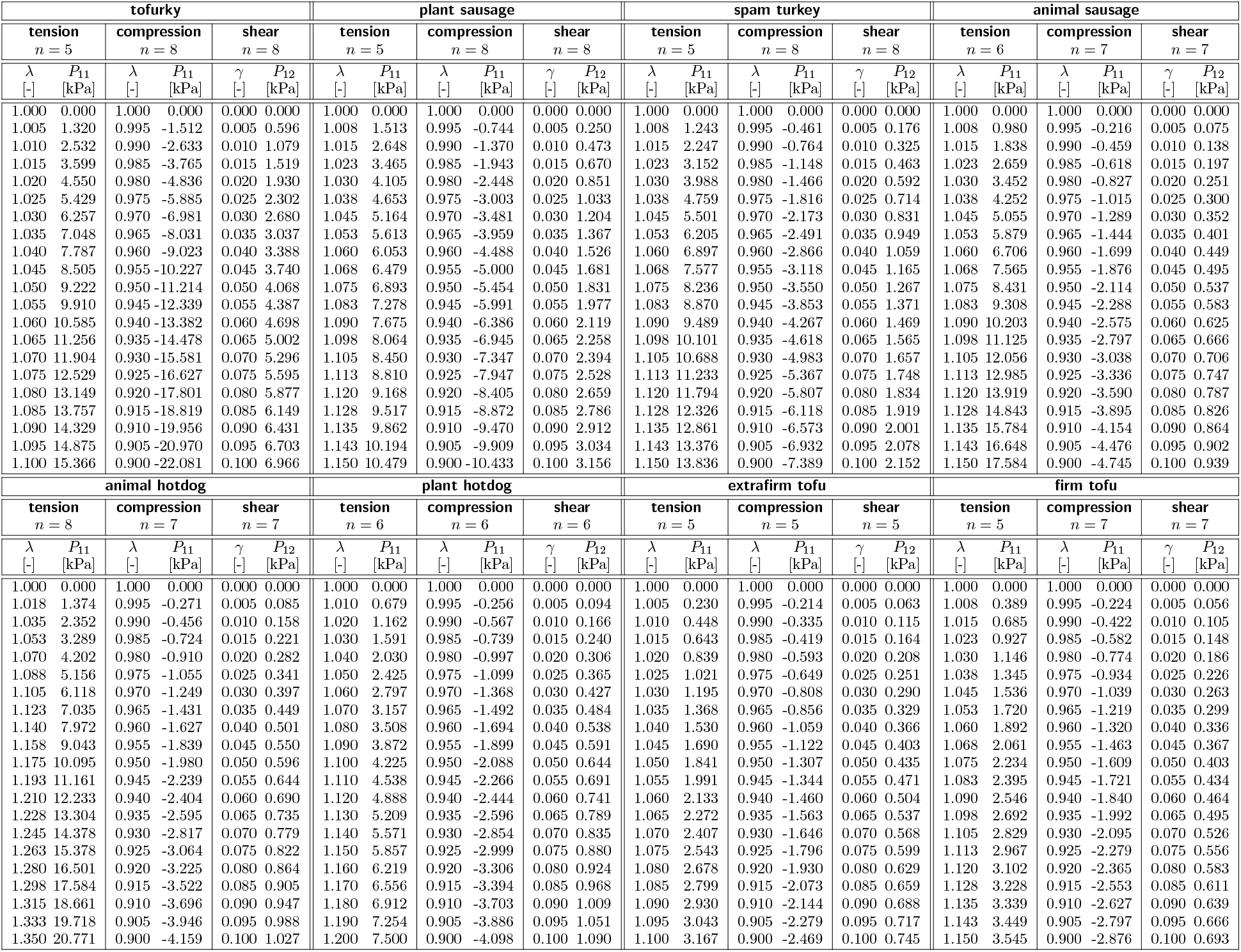
Plant-based and animal meats tested in tension, compression, and shear. Stresses are reported as means from the loading and unloading curves of *n* samples tested in the ranges 1.0 ≤ *λ* ≤ *λ*_max_ for tension, 1.0 ≥ *λ* ≥ 0.9 for compression 0.0 ≤ *γ* ≤ 0.1 for shear. The maximum tensile stretch *λ*_max_ is product dependent and denotes the stretch before failure occurs, so before the stress-stretch curve displays a significant drop.

Figure 1 shows the resulting stress-stretch and shear stress-strain curves with the plant-based products in blue and the animal products in red. The small standard error, highlighted as the shaded region around the mean, reveals that there is little sample-to-sample variation and that our mechanical tests are solidly reproducible. Notably, tofurky displays by far the stiffest response, followed by plant-based sausage, the three animal products, spam turkey, animal sausage, and animal hotdog, and then plant-based hotdog, extrafirm tofu, and firm tofu. Interestingly, the plant-based products, plant-based sausage and hotdog, display similar mechanical properties as their animal counterparts, animal sausage and hotdog, whereas the two tofu products are notably softer.

**Fig. 1.**
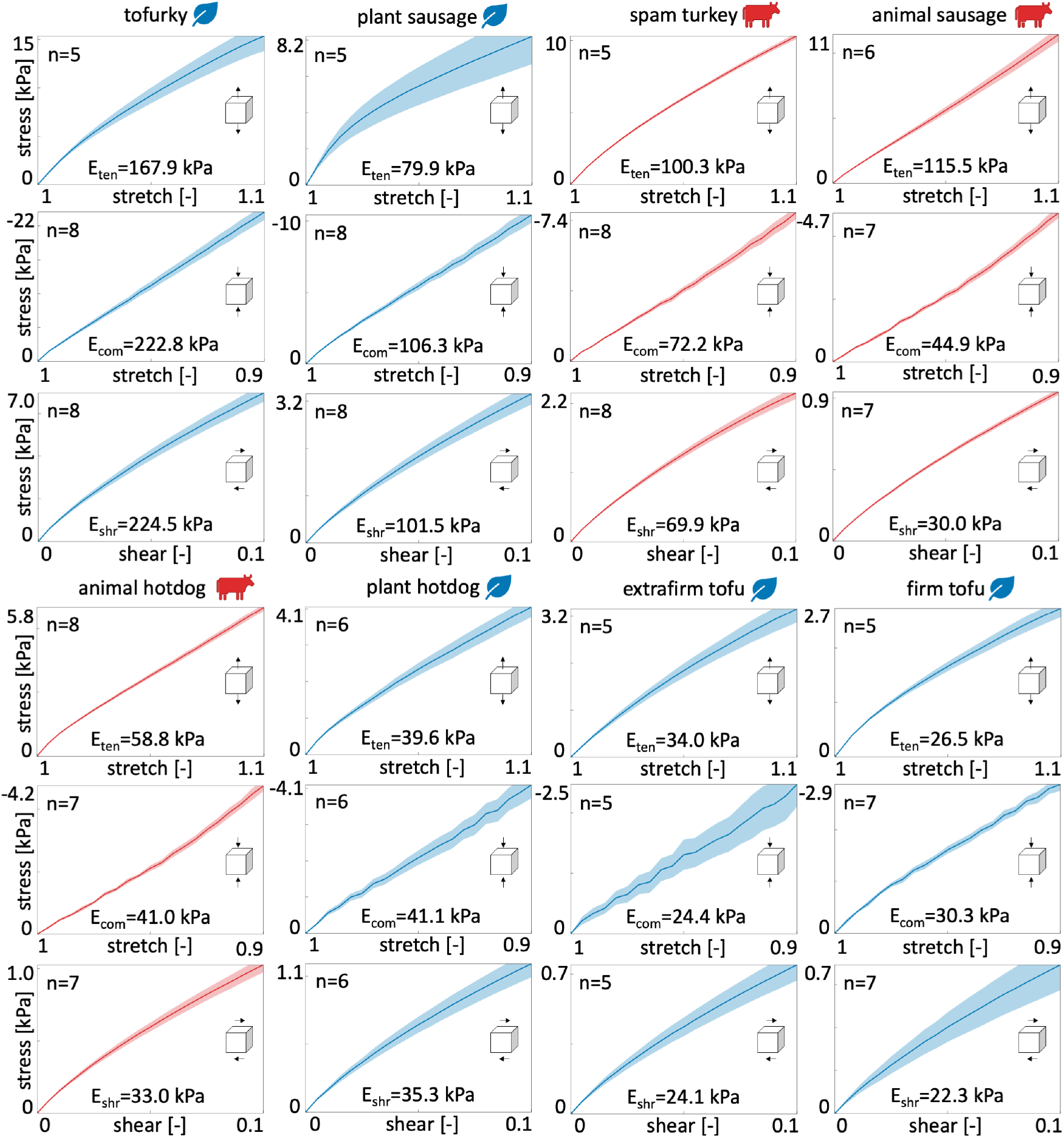
Mechanical testing. Tofurky, plant-based sausage, spam turkey, animal sausage, animal hotdog, plant-based hotdog, extrafirm tofu, and firm tofu tested in tension, compression, and shear. Stresses are reported as means ± standard error of the means of *n* samples tested in the ranges 1.0 ≤ *λ* ≤ 1.1 for tension, 0.9 ≤ *λ* ≤ 1.0 for compression, and 0.0 ≤ *γ* ≤ 0.1 for shear. Each plot reports the sample size *n* and the stiffness, *E*_ten_, *E*_com_, *E*_shr_. Plant-based products are colored in blue, animal products in red.

### Mechanical signatures

We extract the mechanical signatures of all eight products from Table 2 and directly compare the tensile, compressive, and shear stiffnesses, the peak tensile stresses and stretches, and the tension-compression asymmetry of the eight products in Figure 2 and in Table 1. Plant-based products are colored in blue, animal products in red, with darker colors indicating a stiffer response across all testing modes. Figure 2 **a** shows the tensile stress-stretch curves for stretches up to +10%. In addition, we test all tensile samples to failure, where we define failure as a notable drop in the stress response, and plot the tensile stress-stretch curves *before* failure occurs in Figure 2 **b**. Both tofurky and extrafirm tofu exhibit failure slightly above 10% stretch, while all other products stretch up to 15%–20% without failing. Animal hotdog stretches the furthest, up to 35%, and reaches the largest peak stress. Figures 2 **c** and 2 **d** show the compressive stress-stretches curves for stretches up to −10%, and the shear stress-strain curves for shear strains up to 10%. Clearly, across the entire top row, in all three modes, tofurky is the stiffest, and the two tofu products are the softest. From the tension, compression, and shear curves in Figures 2 **b-d**, we extract the linear elastic modulus using linear regression and report it as the tensile, compressive, and shear stiffnesses *E*_ten_, *E*_com_, *E*_shr_ in Figures 2 **e-g**, and as the mean stiffness *E*_mean_ ± standard deviation across all three modes in Figure 2 **h**. Across all three modes, tofurky with a stiffness of 205.1±32.2 kPa is by far the stiffest product. In fact, it is more than twice as stiff as the second and third products, plant-based sausage with 95.9±14.1 kPa and spam turkey with 80.8±16.9 kPa. The two tofu products are consistently the softest products across all three loading modes, where extrafirm tofu with a stiffness of 27.5±5.6 kPa is slightly stiffer than firm tofu with 26.4±4.0 kPa. From the tension-to-failure curves in Figure 2 **b**, we extract the peak tensile stress in Figure 2 **i** and the peak tensile stretch in Figure 2 **j**. Interestingly, animal hotdog displays the largest peak stress and stretch. Tofurky and animal sausage both have high peak stresses, but much lower peak stretches. extrafirm and firm tofu have by far the lowest peak stresses. Figure 2 **k** shows the tension-compression asymmetry at ±10% stretch. The black line indicates tension-compression symmetry; a value larger than one means that the product is stiffer in tension than in compression, and a value smaller than one means the opposite. Strikingly, all three animal products display a larger stiffness in tension than in compression. Animal sausage shows the largest asymmetry with 2.41, followed by animal hotdog with 1.41 and spam turkey with 1.39. The only plant-based product with a comparable tension-compression asymmetry is extrafirm tofu with 1.29. Plant-based hotdog with 1.03, firm tofu with 0.95, plant-based sausage with 0.79 and tofurky with 0.70, all display either close to no asymmetry or a larger stiffness in compression than in tension.

**Fig. 2.**
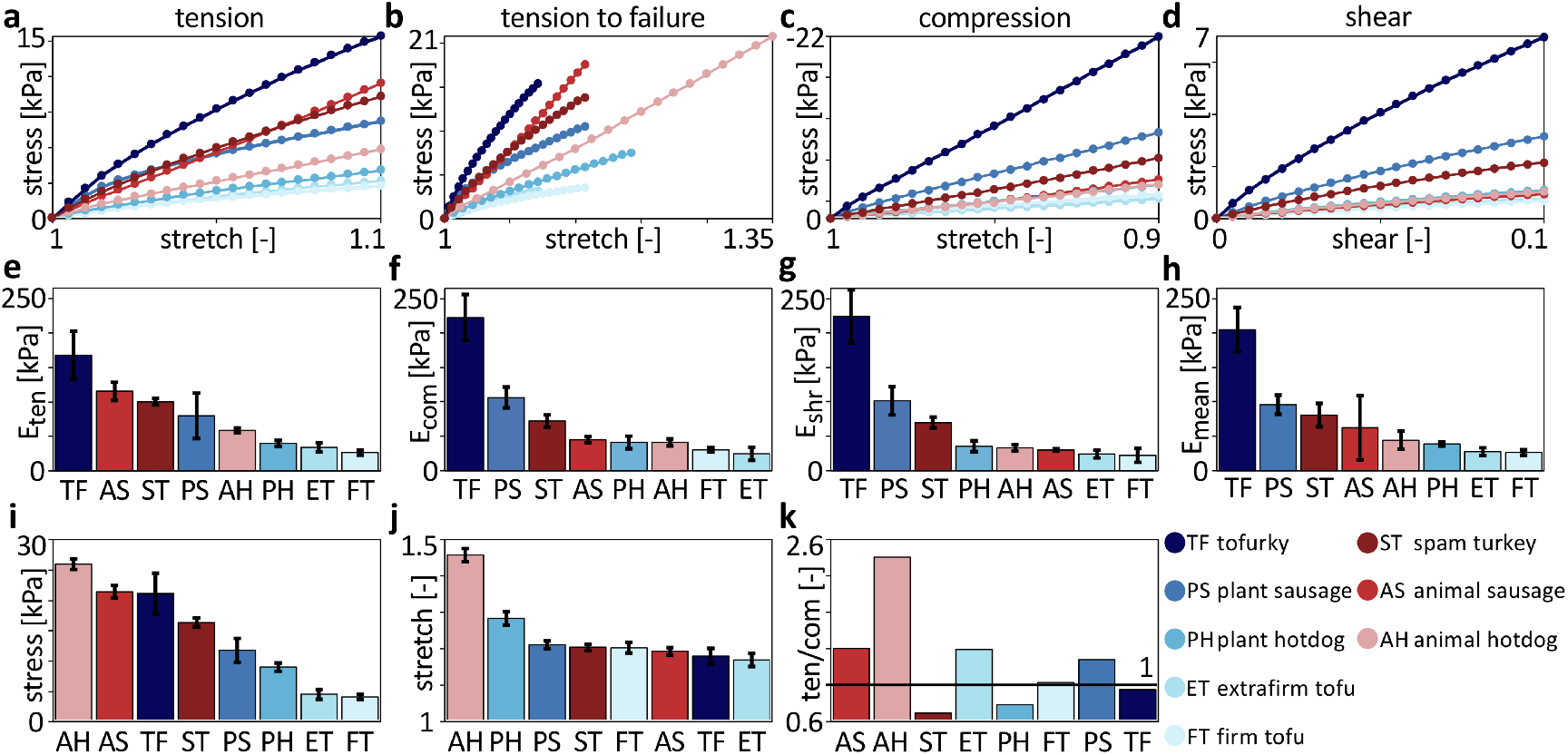
Mechanical signatures. Tofurky, plant-based sausage, spam turkey, animal sausage, animal hotdog, plant-based hotdog, extrafirm tofu, and firm tofu characterized in tension, compression, and shear. **a**. Tensile stress-stretch curves up to +10% stretch. **b**. Compressive stress-stretch curves up to − 10% stretch. **c**. Shear stress-strain curves up to 10% shear strain. **d**. Tensile stress-stretch curves up to failure. **e**. Tensile stiffness *E*_ten_. **f**. Compressive stiffness *E*_com_. **g**. Shear stiffness *E*_shear_. **h**. Mean stiffness *E*_mean_ with standard deviation. **i**. Peak tensile stress, mean and standard deviation. **j**. Peak tensile stretch, mean and standard deviation. **k**. Tension-compression asymmetry in stress at 10% stretch. The black line indicates tension-compression symmetry, products above the line have a greater tensile stress and products below the line have a greater compressive stress. The legend shows the two-letter keys and colors that correspond to each product with plant-based products in blue and animal products in red.

### Model discovery

The mechanical signatures in Figure 2 provide valuable first insight into the material behavior of plant-based and animal meats. However, these signatures are only one-dimensional, and cannot predict the complex material behavior of the eight products in real three-dimensional chewing. We now discover the fully three-dimensional best-in-class one- and two-term models for all eight products using the constitutive neural network in Figure 8.

Figure 3 summarizes the results for the eight different meat products in terms of the best one- and two-term models made up of eight functional building blocks: linear, exponential linear, quadratic, and exponential quadratic in the first and second strain invariants *I*_1_ and *I*_2_. The color code indicates the quality of fit to the tension, compression, and shear data from Table 2, with dark blue indicating the best fit and dark red the worst. Motivated by the notable differences in the discovered models for tensile stretches up to 10% versus 35% in Figure 9, we decide to use the full tension data, not just the 10% stretch. We observe that, with the entire tension regime, the error plot becomes more non-uniform, and the best-in-class models become easier to delineate.

**Fig. 3.**
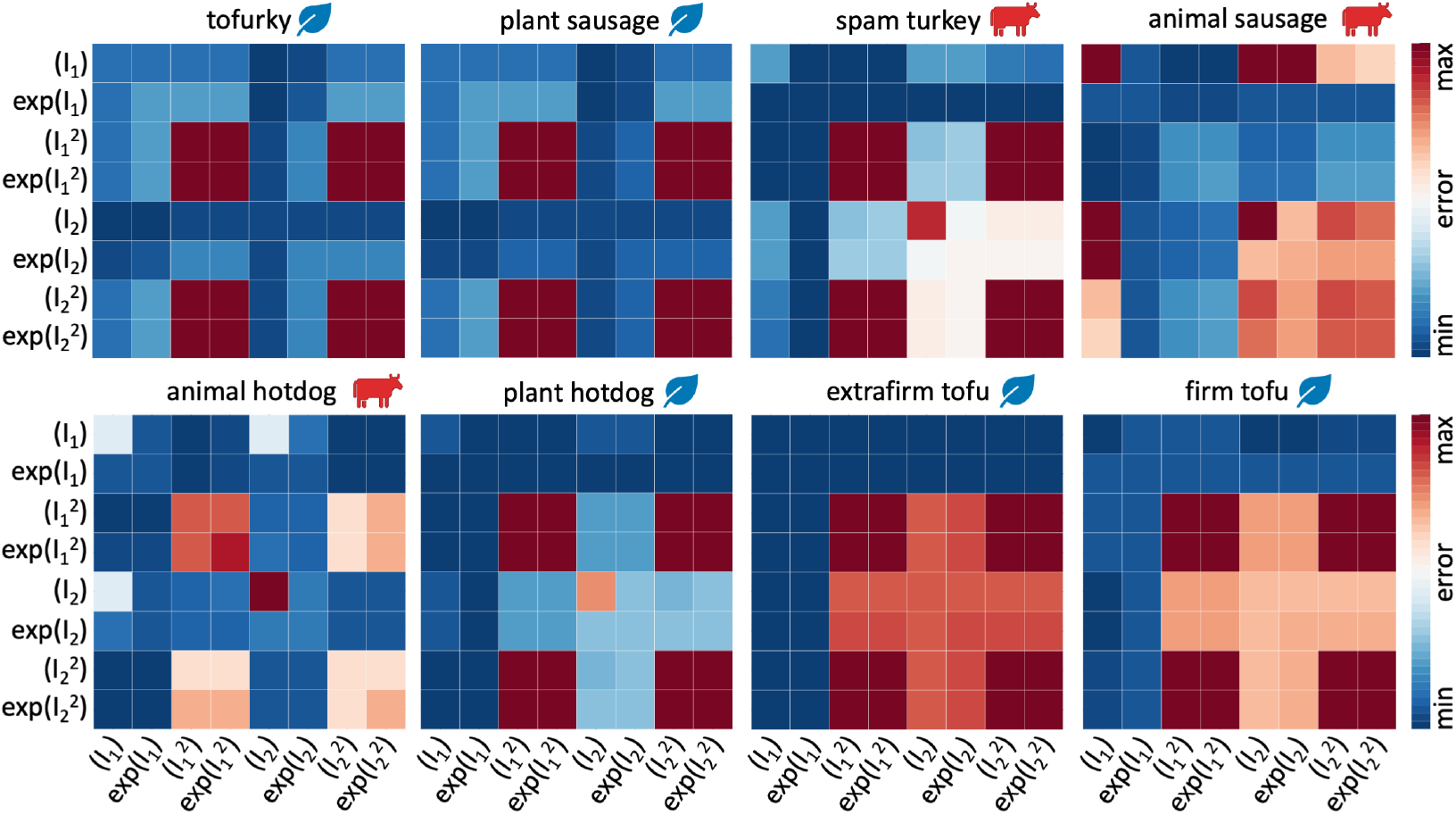
Model discovery. Discovered one-term models, on the diagonal, and two-term models, off-diagonal, for tofurky, plant sausage, spam turkey, animal sausage, animal hotdog, plant hotdog, extrafirm tofu, and firm tofu. All models are made up of eight functional building blocks: linear, exponential linear, quadratic, and exponential quadratic terms of the first and second strain invariants *I*_1_ and *I*_2_. The color code indicates the quality of fit to the tension, compression, and shear data from Table 2, ranging from dark blue, best fit, to dark red, worst fit.

For the best-in-class one-term models that correspond to the bluest most term on the diagonals of Figure 3, we discover three prominent soft matter models: the linear second invariant Blatz Ko model [30], *w*_5_[*I*_2_ - 3], for tofurky and plant-based sausage, the exponential linear first invariant Demiray model [31], 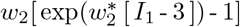, for spam turkey, animal sausage, animal hotdog, and plant-based hotdog, and the linear first invariant neo Hooke model [32], *w*_1_[*I*_1_ - 3], for extrafirm tofu and firm tofu, with the following best-fit parameters,

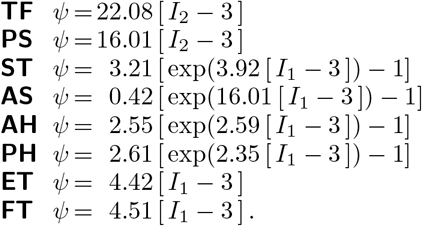

Notably, none of the quadratic terms, 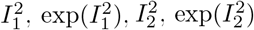, describe the material behavior well, as we conclude from the red squares on the eight diagonals in Figure 3.

For the best-in-class two-term models that correspond to the bluest most term overall, we discover the linear first and second invariant Mooney Rivlin model [33, 34], *w*_1_[*I*_1_ - 3] + *w*_5_[*I*_2_ - 3], for tofurky and plant-based sausage, the linear and quadratic first invariant model, *w*_1_[*I*_1_ - 3] + *w*_3_[*I*_1_ - 3]^2^, for spam turkey and plant-based hotdog, the linear and exponential quadratic first invariant model, *w*_1_[*I*_1_ - 3] + *w*_4_[exp(*w*_4*∗*_ [*I*_2_ - 3]^2^) - 1], for animal sausage, the linear first and quadratic second invariant model, 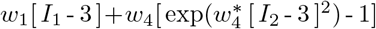, for animal hotdog, and the linear first invariant neo Hooke model [32], *w*_1_[*I*_1_ - 3], for extrafirm tofu and firm tofu, with the following best-fit parameters,

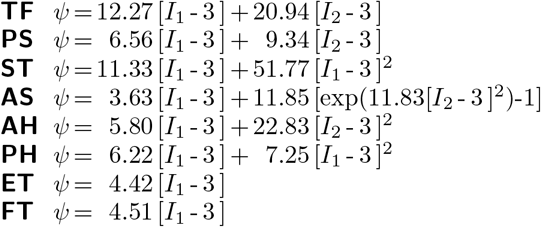

Notably, all eight models contain the classical linear first invariant neo Hooke term [32], [*I*_1_ −3]. All eight products, except for animal sausage, show the same four dark red corners patterns, which indicate that the 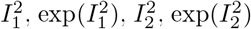 terms provide the worst fit to the data. From the nearly identical error maps, we conclude that tofurky and plant-based sausage have a similar mechanical behavior, and so do plant-based hotdog, extrafirm tofu, and firm tofu. In contrast, the three animal products, spam turkey, animal sausage, and animal hotdog, have the most distinct error maps, suggesting that their mechanical behavior is notably different. Interestingly, for both tofu products, the model fit does not improve by adding a second term, and their best-in-class two-term model is identical to their best-in-class one-term model: the classical neo Hooke model [32]. To visualize how well these discovered models approximate to our mechanical tests, we color-code the stress contributions of the individual terms and illustrate them together with the raw data in Figure 4. The eight columns correspond to each of the eight products, and the rows represent the tension, compression, and shear experiments. The dark red term that consistently appears in all eight models is the classical linear first invariant neo Hooke term [32], *w*_1_ [*I*_1_ − 3]. In addition, both tofurky and plant-based sausage have a turqouise linear second invariant Blatz Ko term [30], *w*_5_ [*I*_2_− 3], spam turkey and plant-based hotdog have an orange quadratic first invariant term, *w*_3_ [*I*_1_− 3]^2^, animal sausage has a yellow exponential quadratic first invariant term, *w*_4_ exp 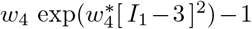, animal hotdog has a blue quadratic second invariant term, *w*_7_ [*I*_2_− 3]^2^, and extrafirm tofu and firm tofu have no additional second term.

**Fig. 4.**
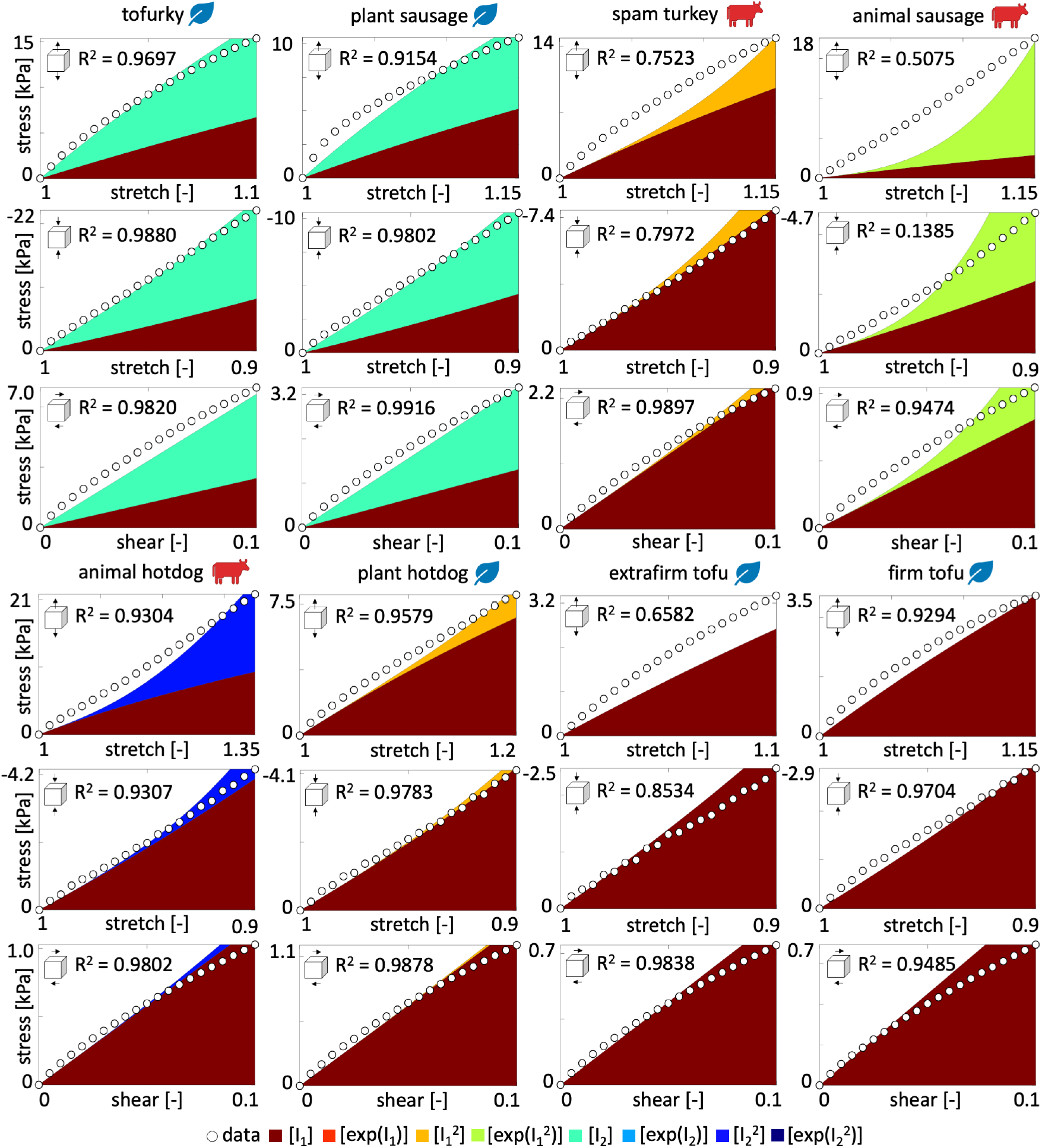
Stress as a function of stretch or shear strain for the best-in-class two-term model. We train the constitutive neural network in Figure 8 simultaneously with the tension, compression, and shear data from Table 2, and apply *L*_0_-regularization to reduce the number of terms to two according to Table 1 and Figure 3. The color-coded regions designate the contributions of the eight model terms to the stress function according to Figure 8. The coefficients of determination R^2^ indicate the goodness of fit.

Table 1 summarizes the discovered best-in-class one- and two-term models and parameters. For the Mooney Rivlin type models, we can directly translate the network weights into the shear modulus *µ* using *µ* = 2 *w*_1_ + 2 *w*_5_. So, the shear moduli of the two stiffest products, tofurky and plant-based sausage, are 66.42 kPa and 31.80 kPa, and the shear moduli of the two softest products, extrafirm and firm tofu, are 8.84 kPa and 9.04 kPa. Interestingly, the shear modulus for firm tofu is slightly larger than for extrafirm tofu. We can translate these shear moduli *µ* into Young’s moduli, *E* = 2 [1 + *ν*] *µ*, by assuming a Poisson’s ratio of *ν* = 0.5 for perfect incompressibility, and we find Young’s moduli of 199.25 kPa, 95.40 kPa, 26.53 kPa, and 27.08 kPa for tofurky, plant-based sausage, extrafirm tofu, and firm tofu. These values are in excellent agreement with the mean stiffnesses from our linear regression in Figure 2 **h**, 205.09 kPa, 95.89 kPa, 27.48 kPa, and 26.35 kPa for tofurky, plant-based sausage, extrafirm tofu, and firm tofu. For the other four products, spam turkey, animal sausage, animal hotdog, and plant-based hotdog, we discover novel constitutive models with quadratic or exponential terms that do not directly translate into Young’s moduli. Interestingly, all three animal products are in this group, indicating that animal meat, even when highly processed, has a more complex mechanical behavior than plant-based meat.

### Food texture survey

To explore to which extent our perception of taste aligns with our discovered mechanical signatures, we survey *n* = 16 participants to characterize the textural properties of our eight products after eating a small sample. Before the texture survey, all participants perform two baseline surveys: the Food Neophobia Survey, which probes how open participants are to trying new foods [35], and the Meat Attachment Questionnaire, which probes how attached participants are to eating meat [36]. From the results in Figure 10, we conclude that our participants are very open to trying new food and are ambivalent to having meat in their diet. All participants then rank each meat product on a 5-point Likert scale, ranging from 5 for strongly agree to 1 for strongly disagree, based on how well each sample agrees with twelve traditional texture features [21, 22]. Figure 5 summarizes the results for all twelve features, with the eight products ordered by the level that participants agreed to most. Blue colors indicate plant-based products and red colors indicate animal products. Interestingly, six of the twelve texture features, *brittle, gummy, viscous, springy, sticky*, and *fibrous*, display variations that are all well below 1.3, and not statistically significant across the eight products. This suggests that there is no perceived difference between plant-based and animal products in these six texture features. The other six texture features, *soft, hard, chewy, moist, fatty*, and *meat-like*, display significant variations, all but *chewy* well above 2.5 across the eight products. In agreement with our intuition, *soft* and *hard* are perceived as inversely correlated, with a Spearman rank correlation coefficient of *ρ* = −0.9286 and *p* = 0.0022, with the first, second, third, fourth, and eighth products on the softness scale, firm tofu at 4.8±0.4, extrafirm tofu at 4.5±0.6, spam turkey at 4.3±0.6, animal hotdog at 3.7±1.0, and tofurky at 1.7±0.9, scoring inversely on the hardness scale and the fifth, sixth, and sevenths products, animal sausage at 3.4±1.1, plant-based hotdog at 3.4±0.9, and plant-based sausage at 3.1±1.0, scoring almost identically for both, soft and hard. Strikingly, *fatty* and *meat-like* are perceived as correlated, with a Spearman rank correlation coefficient of *ρ* = 0.9762 and *p* = 0.0003, with the three animal products, animal sausage at 4.4±1.1 and 4.9±0.3, animal hotdog at 3.8±1.0 and 3.9±1.3, and spam turkey at 3.6±1.1 and 3.7±0.4, scoring first, second, and third in both categories. The dominance of animal sausage is interesting, as it is the least complex of the animal products. The three plant-based products, plant-based hot-dog, plant-based sausage, and tofurky, score fourth, fifths, and sixth. Naturally, the two tofu products, extrafirm tofu at 1.9±1.1 and 1.3±0.6 and firm tofu at 1.5±0.8 and 1.2±0.4, score by far the lowest in the plant-based category. The animal meats are the three highest ranked for fattiness with plant-based sausage and plant-based hotdog next. Both tofu products and tofurky rank much lower. All products rank fairly highly for moistness, except for plant-based sausage and tofurky. Most notably, the food tasting survey produces nearly identical stiffness rankings as the mechanical testing, with a Spearman rank correlation coefficient of *ρ* =0.8333 and p=0.0154.

**Fig. 5.**
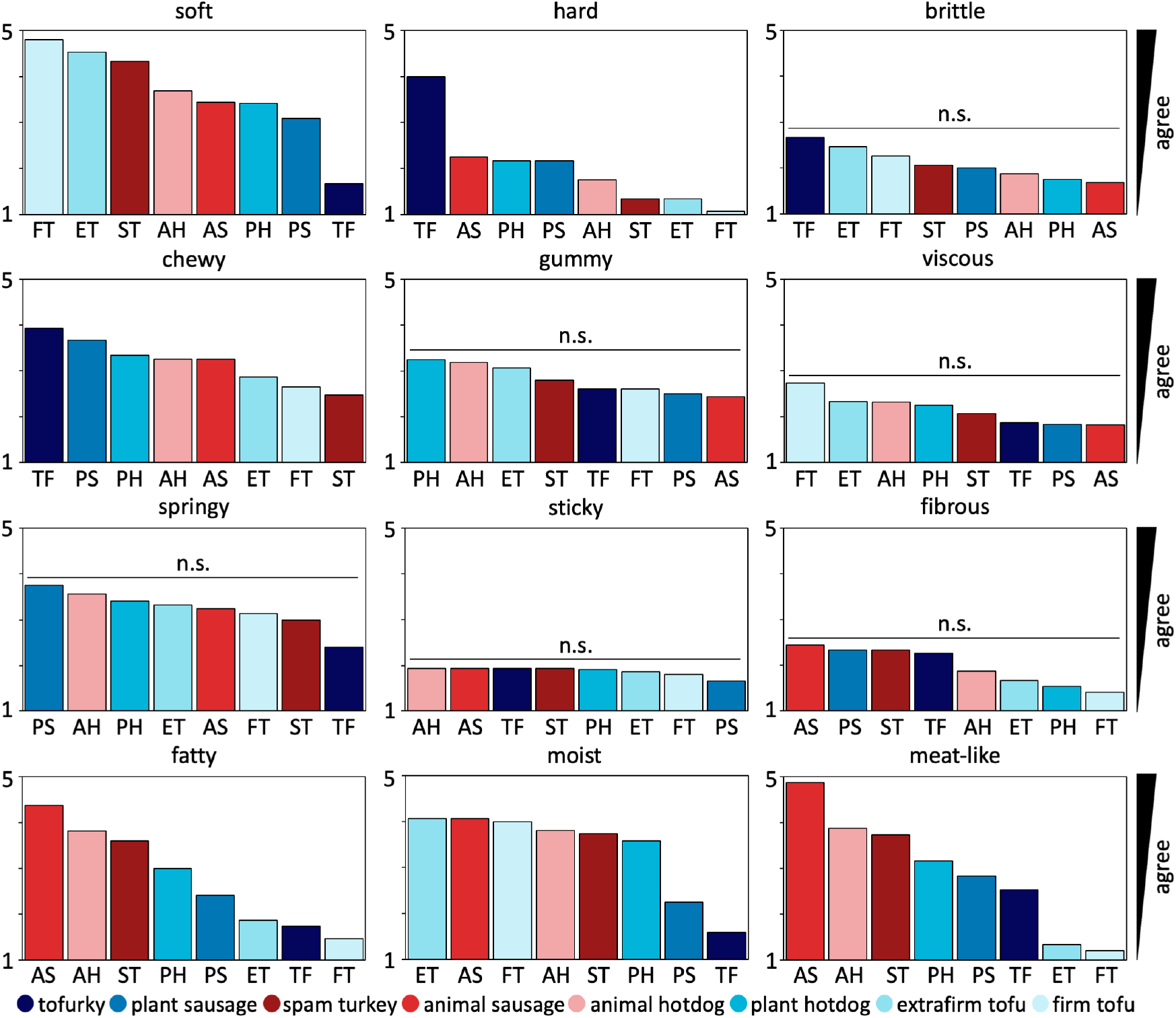
Food texture survey. Tofurky, plant sausage, spam turkey, animal sausage, animal hotdog, plant hotdog, extrafirm tofu, and firm tofu profiled for twelve texture features. Participants eat cooked samples of all eight products, and rank their texture feature on a 5-point Likert scale ranging from 5 for strongly agree to 1 for strongly disagree. Each survey question asks “this food is [texture feature]” with texture features *soft, hard, brittle, chewy, gummy, viscous, springy, sticky, fibrous, fatty, moist*, and *meat-like*, adopted from traditional texture classifications [21, 22]. Within each plot, meat products are sorted from highest to lowest agreement, from left to right. Plant-based meats are colored in blue, animal meats in red. n.s. denotes that variations are not statistically significant.

## Discussion

We tested five plant-based products, tofurky, plant-based sausage, plant-based hotdog, extrafirm and firm tofu, and three animal products, spam turkey, animal sausage, and animal hotdog, in tension, compression, and shear. We focused on highly processed plant-based and animal meats, assuming that all products are nearly isotropic and homogeneous, and, as our study confirms, easy and reproducibly to test. We performed a total of 157 mechanical tests and 288 neural network simulations. Our goal was to probe: *To which extent do plant-based meat products mimic the mechanical signature of animal meat?*, not just in a double-compression texture profiling analysis, but across the entire three-dimensional spectrum. While our study is limited by our assumption of isotropy and by probing raw products at room temperature, it uncovers several interesting and unexpected results:

### Plant-based sausage and hotdog succeed in mimicking the mechanical signature of their animal counterparts

Our results confirm that our mechanical tension, compression, and shear tests are well reproducible with narrow errors and standard deviations, as we conclude from Figure 1 and Table 1. From the mean tension, compression, and shear curves in Figures 2 **a,b,c,d**, we conclude that plant-based sausage and hotdog consistently lie in the middle range of all eight products, and place closer to the three animal meats than the two tofu products. When comparing at the stiffnesses in Figures 2 **e,f,g,h**, plant-based sausage and hotdog range from *E* = 35.3 kPa to *E* = 106.3 kPa and fall in a comparable category as animal sausage and hot-dog ranging from *E* = 26.8 kPa to *E* = 115.5 kPa. In contrast, tofurky ranges from *E* = 167.9 kPa to *E* = 224.5 kPa and is consistently more than twice as stiff, while the two tofu products range from *E* = 22.3 kPa to *E* = 34.0 kPa and are about half as stiff. Our discovered stiffness values lie well within the range of the reported compression stiffnesses for sausage of *E* = 120 kPa, turkey of *E* = 90 kPa, and chicken of *E* = 40 kPa [25] using traditional texture profiling analysis [37]; the reported tensile stiffnesses for chicken of *E* = 360 kPa and soy protein of *E* = 100 kPa using tensile testing [26]; and our previous study of tofurky of *E* = 282 kPa, plant-based chicken of *E* = 108 kPa, and real chicken of *E* = 87 kPa using tension, compression, and shear testing [28]. Our results suggests that, when looking for vegetarian options, we no longer have to rely on tofu alone. Tofu, a protein-and vitamin-rich product of curdled soy milk, was first produced nearly 2,000 years ago in China, and has since then become a popular substitute for meat worldwide [38]. Today, tofu comes in various stiffnesses: soft, silken, regular, firm, and extrafirm. Its stiffness is tunable by its water content that ranges from 85-90% for soft to 40%-50% for extrafirm. In our study, in Table 1 and Figures 1 and 2, even firm and extrafirm tofu consistently display the lowest stiffness across all products and tests, and perform poorly in reproducing the mechanical properties of animal meat. In contrast, plant-based sausage and hotdog successfully mimic the mechanical signature of their animal counterparts. Interestingly, market analyses predict that, of all different plant-based meat products, burgers, patties, ground, nuggets, and sausages, the sausage market will experience the highest growth rate within the next decade [39]. As one of the most common protein-based breakfast foods, plant-based sausage comes in a variety of flavors and is in high demand worldwide. Yet, as we can see in Table 1, the growing success of plant-based sausage and hotdog comes at a price: a high sodium content, added sugars, and long list of additives and ingredients [40].

### The more complex the product, the more complex its mechanics

In this study, we perform the first fully three-dimensional characterization of eight different plant-based and animal meat products. We analyze the data using a constitutive neural network with *L*_0_-regularization [29] to discover the best one- and two-term models that simultaneously fit the tension, compression, and shear data for each product. Using the error plots in Figure 3 and the discovered weights in Table 1, we can easily write out the free energy functions that best fit each meat. Strikingly, the oldest and simplest of all models, the classical widely used neo Hooke model [32] in terms of only *I*_1_ is the best model for firm and extrafirm tofu, the two softest, oldest, and simplest products with the shortest list of ingredients: water, soybeans, calcium sulfate, and calcium or magnesium chloride. The popular Mooney Rivlin model [33, 34] in terms of *I*_1_ and *I*_2_ is the best model for tofurky and plant-based sausage, the two stiffest products. Interestingly, we discover three novel, nonlinear material models for the three highly processed animal meats: animal sausage in terms of *I*_1_ and 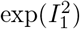, animal hotdog in terms of *I*_1_ and 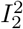, and spam turkey in terms of *I*_1_ and 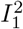. This suggests that processed animal meat has a complex mechanical behavior that is not appropriately acknowledged by common existing constitutive models. Surprisingly, the degree of complexity of our discovered material models, from one-term to two-term and from linear to quadratic to exponential, mimics the complexity of the ingredient list, from a few pure ingredients for tofu to a wide of variety additives and ingredients for sausage and hotdog. Discovering product-specific best-fit models from data would have been unthinkable one or two decades ago, and is only now made possible by recent developments in constitutive neural networks, machine learning, and artificial intelligence [29]. This suggests that, instead of using a trial-and-error approach to improve the texture of plant-based meat, we could envision using generative artificial intelligence to scientifically generate recipes for plant-based meat products with precisely desired properties.

### Animal products are stiffer in tension than in compression, while plant-based products are not

An insightful mechanical property that is impossible to quantify by a double-compression texture profiling analysis alone [24] is the tension-compression asymmetry. Our combined tension, compression, and shear tests in Figures 1 and 2 **k** reveal that the three animal meats display the highest tension-compression asymmetry, with 2.41 for animal sausage, 1.41 for animal hotdog, and 1.39 for spam turkey. Interestingly, all plant-based products, except for extrafirm tofu, are either close to symmetric or stiffer in compression than in tension with values smaller than one, with 1.03 for plant-based hotdog, 0.95 for firm tofu, 0.79 for plant-based sausage, and 0.70 for tofurky. Similarly, all three animal meats rank amongst the four products with the highest failure stress of 26.0 kPa for animal hotdog, 21.4 kPa for animal sausage, and 16.3 kPa for spam turkey, in 2 **i**, while all plant-based products, except for tofurky, have significantly lower failure stresses. Understanding the mechanical properties of plant-based protein products is a rapidly growing field of research [26]. Naturally, the tensile, compressive, and shear stiffnesses in Figure 2 are highly sensitive to plant source and processing [37], and successful formulations often benefit from protein synergies, such as soy and wheat or pea and potato [14]. In addition, the complex ingredient lists in Table 1 suggest that tunability may require other non-protein components such as oil or starch. One known short-coming of plant-based meats is that they lack adipose tissue, which contributes to the mouthfeel, appearance, and texture of meat [41], and these additives could address this limitation. The quest for finding the best ingredients raises an interesting question: Instead of applying our technology only to a *forward analysis*, where we test an existing product and characterize its mechanical features, can we perform an *inverse analysis*, where we prescribe desired mechanical features and determine the required ingredient list? Our study demonstrates that constitutive neural networks provide a powerful tool to learn functional mappings between products and texture [28]. What if we could expand this technology to learn inverse mappings between texture and ingredients to fine-tune the mechanical signature of plant-based meat?

### Our perception of stiffness matches mechanical testing

As we conclude from the food texture survey in Figure 5, the rankings for softness and hardness closely match the ranking for stiffness in Figure 2 **h**: Tofurky ranks the least soft, the hardest, and the stiffest, while firm and extrafirm tofu rank the softest, the least hard, and the least stiff. The results of our survey suggest that some textural features are easier for the participants to delineate than others. For example, the participants perceived significant differences in soft, hard, chewy, fatty, moist, and meat-like across all eight products, but were unable to delineate products by brittle, gummy, viscous, springy, sticky, and fibrous. The lack of variation in these six features can be explained, at least in part, by a similar perception across all eight products, or by a lack of clarity of their interpretation.

In summary, our results confirm that in seeking to design plant-based alternatives that truly mimic the sensory experience of animal meat, it is not sufficient to solely rely on traditional one-dimensional double compression tests. Instead, we suggest to consider fully three-dimensional testing to discover the true material behavior across a broad loading range, and understand the subtle mechanical differences that may trigger differences in our perception of taste. Our study shows that the one-dimensional stiffness of plant-based and animal sausage and hotdog is nearly identical, but their three-dimensional characteristics are not. Our approach to automatically discover the mechanics of plant-based and animal meat with constitutive neural networks could be a starting point towards using generative artificial intelligence to reverse-engineer formulas for plant-based meat products with customer-friendly tunable properties. We hope that the present study encourages others, especially in academia or nonprofit organizations who can freely share their results, to undertake complementary studies and contribute to an open source data base to accelerate discovery and innovation towards a more efficient and sustainable global food system.

## Methods

### Mechanical testing

We test five plant-based meat products: ham style roast tofurky (Tofurky, Hood River, OR), artesian vegan frankfurter plant sausage (Field Roast, Seattle, WA), signature stadium plant hotdog (Field Roast, Seattle, WA), organic firm tofu (365 by Whole Foods Market, Austin TX), organic extrafirm tofu (House Foods, Garden Grove, CA). For comparison, we also tested three animal meat products: wieners classic animal hotdog (Oscar Meyer, Kraft Heinz Co, Chicago, IL), oven roasted spam turkey (Spam, Hormel Foods Co, Austin, MN), and turkey polska kielbasa animal sausage (Hillshire Farm, New London, WI). Table 1 summarizes the ingredients of all eight products. For each meat type, we test at least *n* = 5 samples in tension, compression, and shear. Figure 6 documents our sample preparation and our mechanical testing.

**Fig. 6.**
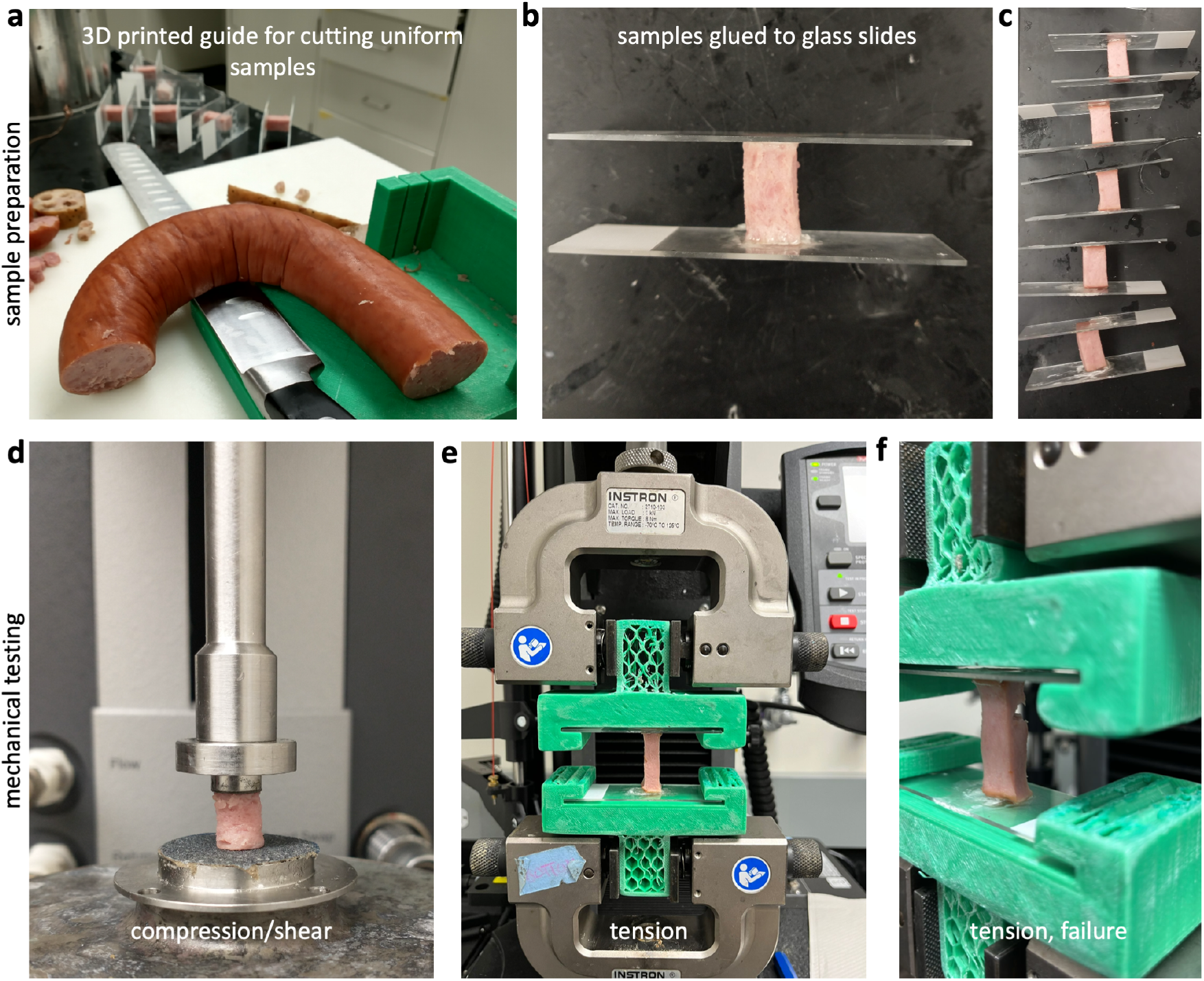
Sample preparation and mechanical testing. **a**. Samples are cut using a 3D printed guide and a brain sectioning knife to obtain uniform dimensions of 1 ×1 ×2 cm^3^ for tensile testing. **b**. Samples are super glued to glass microscope slides. **c**. Samples are left to set for 30-45 minutes for the glue to adhere. **d**. Cylindrical samples of 8 mm diameter and 1 cm height are cored using a biopsy punch and loaded into a rheometer for compression and shear testing using parallel plates with sandpaper on both surfaces. **e**. Tension samples are loaded into 3D printed grips, which are compatible with an Instron testing device. **f**. Tension test is run until the sample fails. **a, b, d** show animal sausage and **c, e, f** show animal hotdog.

### Sample preparation

We prepare the samples following our established protocols [28]. Figures 6 **b,c,e,f** illustrate our tension tests, for which we use a custom 3D printed cutting guide and brain sectioning knife to prepare samples of 1 ×1 ×2 cm^3^. We super glue the samples to glass microscope slides and wait for 30 minutes until the glue is fully cured. During curing, we drape a damp paper towel over the samples to keep them hydrated. For the compression and shear tests, we prepare cylindrical samples of 8 mm diameter and 1 cm height using a biopsy punch to extract full-thickness cores from the center of each material, and store the samples in a damp paper tower until testing.

### Sample testing

For all three modes, tension, compression, and shear, we test the samples raw and at room temperature at 25°C [28]. We perform all uniaxial tension tests using an Instron 5848 (Instron, Canton, MA) with a 100N load cell, see Figure 6 **e**. We use 3D printed custom grips to rapidly mount and unmount the microscope slides for high throughput testing. Figure 7 shows the part dimensions to create these grips. We mount each sample in the grips, apply a small pre-load, and calibrate the initial gauge length *L*. We determine the pre-load magnitude for each sample individually. We define pre-load as the minimum load needed to remove any slack, based on visual inspection of a force-displacement curve starting with a fully unloaded specimen, generally on the order of 0.5 N. We then increase the tensile stretch quasi-statically at a rate of 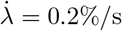, until the sample fails. We perform all uniaxial compression and shear tests using an AR-2000ex torsional rheometer (TA Instruments, New Castle, DE), see Figure 6 **d**. For the compression tests, we mount the sample, apply a small pre-load, and calibrate the initial gage length *L*. We determine the pre-load for each sample based on its loading curve, with values on the order of 0.5 N. We then increase the compressive stretch quasi-statically at a rate of 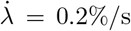 for *t* = 50 s, up to a total stretch of *λ* = 0.9. For the shear tests, we apply a 10% compressive pre-load and calibrate the initial gage length *L*. We then rotate the upper plate quasi-statically at a shear rate of 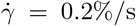 for *t* = 50 s, up to a total shear of *γ* = 0.1. To prevent slippage of the samples during the shear tests, we use a sandpaper-covered base plate of 20 mm diameter and a sandpaper-covered top plate of 8 mm diameter.

**Fig. 7.**
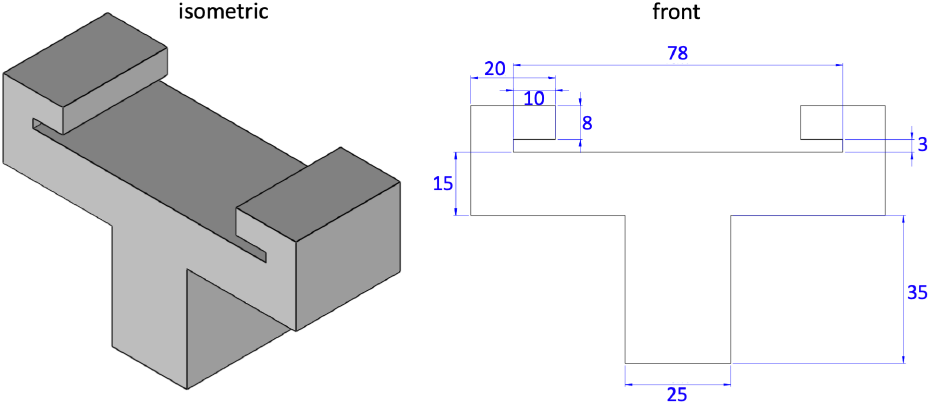
Design of mechanical device for tensile testing. Front, right, and isometric views of our customized 3D printed device to hold glass slides during tensile testing. Two holders need to be printed to complete the set-up. A standard microscope glass slide fits into the 3 mm slot; it mounts and unmounts easily for high throughout testing. All dimensions are in mm.

### Analytical methods and data processing

For each sample and each test mode, we use MATLAB (Mathworks, Natick, MA, USA) to smooth the curves using `smoothingspline` and `SmoothingParam = 1`. We interpolate each smooth curve over 21 equidistant points in the ranges 1.0 ≤ *λ* ≤ *λ*_max_ for tension, 1.0 ≥ *λ* ≥ 0.9 for compression, and 0.0 ≤ *γ* ≤ 1.0 for shear. For each meat product, we select *λ*_max_ in tension as the maximum stretch in the hyperelastic regime, the loading range within which we observe no visible failure for any of the samples. Finally, we average the interpolated curves to obtain the mean and standard error of the mean for each product and report the data in Table 2.

### Stiffness

For each testing mode, we extract the stiffness of each product from the data in Table 2 using linear regression. We convert that the tension and compression data {*λ*; *P*_11_} into strain-stress pairs {*ε*; *σ*}, where *ε* = *λ* − 1 is the strain and *σ* = *P*_11_ is the stress. We postulate a linear stress-strain relation, *σ* = *E · ε*, and use linear regression to determine the tensile and compressive stiffnesses 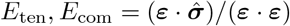. Similarly, we rewrite the shear data, {*γ*; *P*_12_}, as shear strain-stress pairs {*γ*; *τ*}, where *γ* is the shear strain and *τ* = *P*_12_ is the shear stress. We postulate a linear shear stress-strain relation, *τ* = *µ · γ*, convert the shear modulus *µ* into the shear stiffness, *E*_shr_ = 2[1 + *ν*] *µ* = 3 *µ*, and use linear regression, to determine the shear stiffness 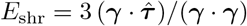.

### Kinematics

We analyze all three testing modes combined using finite deformation continuum mechanics [42, 43]. During testing, particles ***X*** of the undeformed sample map to particles, ***x*** = ***φ***(***X***), of the deformed sample via the deformation map ***φ***. Similarly, line elements of the d***X*** of the undeformed sample map to line elements, d***x*** = ***F*** *·* d***X***, of the deformed sample via the deformation gradient, 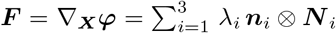. Its spectral representation introduces the principal stretches *λ*_*i*_ and the principal directions ***N*** _*i*_ and ***n***_*i*_ in the undeformed and deformed configurations, where ***F*** *·****N*** _*i*_ = *λ*_*i*_***n***_*i*_. We assume that all meat samples are isotropic and have three principal invariants, 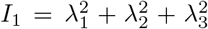 and 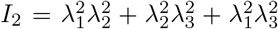 and 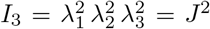, which are linear, quadratic, and cubic in terms of the principal stretches squared. We also assume that all samples are perfectly incompressible, and their third invariant always remains equal to one, *I*_3_ = 1. The remaining two invariants, *I*_1_ and *I*_2_, depend on the type of experiment.

### Constitutive equations

Constitutive equations relate a stress like the Piola or nominal stress ***P***, the force per undeformed area that we measure during our experiments, to a deformation measure like the deformation gradient ***F*** [28]. For a hyperelastic material that satisfies the second law of thermodynamics, we can express the Piola stress, ***P*** = *∂ψ*(***F***)/*∂****F*** − *p* ***F*** ^-t^, as the derivative of the Helmholtz free energy function *ψ*(***F***) with respect to the deformation gradient ***F***, modified by a pressure term, − *p* ***F*** ^-t^, to ensure perfect incompressibility. Here, the hydrostatic pressure, 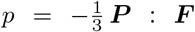, acts as a Lagrange multiplier that that we determine from the boundary conditions of our experiments. Instead of formulating the free energy function directly in terms of the deformation gradient *ψ*(***F***), we can express it in terms of the invariants, *ψ*(*I*_1_, *I*_2_), and obtain the following expression, ***P*** = *∂ψ*/*∂I*_1_ *· ∂I*_1_/*∂****F*** + *∂ψ*/*∂I*_2_ *·∂I*_2_/*∂****F*** − *p* ***F***^-t^.

### Constitutive neural networks

Motivated by these kinematic and constitutive considerations, we reverse-engineer our own constitutive neural network that satisfies the conditions of thermodynamic consistency, material objectivity, material symmetry, incompressibility, constitutive restrictions, and polyconvexity by design [29, 44]. Yet, instead of building these constraints into the loss function, we hardwire them directly into our network input, output, architecture, and activation functions [45, 46] to satisfy the fundamental laws of physics. Special members of this family represent well-known constitutive models, including the neo Hooke [32], Blatz Ko [30], Mooney Rivlin [33, 34], and Demiray [31] models, for which the network weights gain a clear physical interpretation [44, 47]. Specifically, our constitutive neural network learns a free energy function that is parameterized in terms of the first and second invariants. It takes the deformation gradient ***F*** as input, computes the two invariants, *I*_1_ and *I*_2_, raises them to the first and second powers, (○) and (○)^2^, applies either the identity or exponential function, (○) and exp(○), and summarizes all eight terms in the strain energy function *ψ* as the network output. Figure 8 illustrates our network with *n* = 8 nodes, with the following eight-term free energy function,

**Fig. 8.**
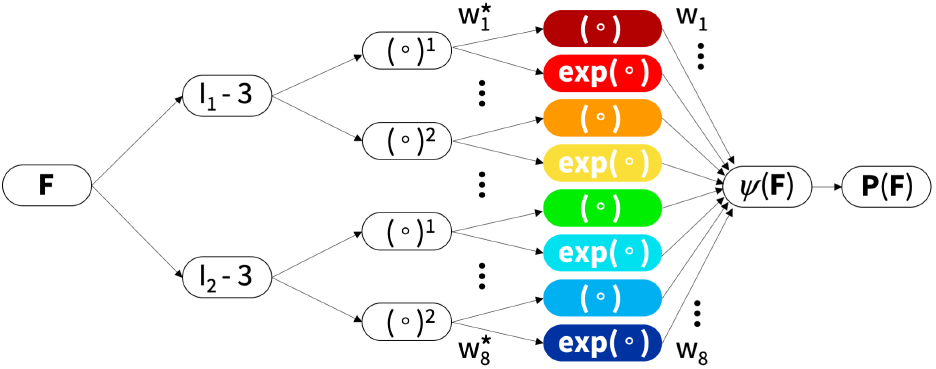
Constitutive neural network. Isotropic, perfectly incompressible constitutive artificial neural network with two hidden layers and eight terms. The network takes the deformation gradient ***F*** as input and calculates its first and second invariant terms, [*I*_1_ − 3] and [*I*_2_ − 3]. The first layer generates powers of these invariants, (○) and (○), and the second layer applies the identity and the exponential function to these powers, (○) and exp(○). The strain energy function *ψ*(***F***) is a sum of the resulting eight terms. Its derivative defines the Piola stress, *∂ψ*(***F***)/*∂****F***, whose components, *P*_11_ or *P*_12_, enter the loss function to minimize the error with respect to the tension, compression, and shear data. By minimizing the loss function, the network trains its weights ***w*** and ***w***^*∗*^ and discovers the best model and parameters to explain the experimental data.

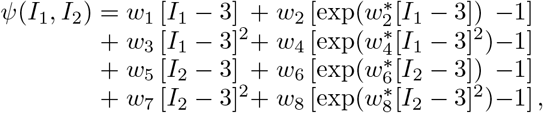

where ***w*** = [*w*_1_, *w*_2_, *w*_3_, *w*_4_, *w*_5_, *w*_6_, *w*_7_, *w*_8_] and 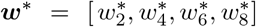 are the network weights. From the derivative of the free energy, we calculate the stress,

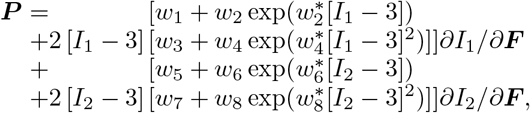

where the derivatives of the invariants, *∂I*_1_/*∂****F*** and *∂I*_2_/*∂****F***, depend on the type or experiment. During training, our network autonomously discovers the best subset of activation functions from 2^8^ − 1 = 255 possible combinations of terms. At the same time, it naturally trains the weights of the less important terms to zero.

### Uniaxial tension and compression

In the tension and compression experiments, we apply a stretch *λ* = *l*/*L*, that we calculate as the ratio between the current and initial sample lengths *l* and *L*. We can write the deformation gradient ***F*** in matrix representation as

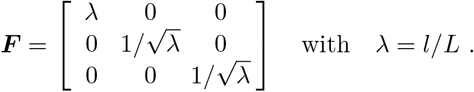

In tension and compression, the first and second invariants and their derivatives are *I*_1_ = *λ*^2^ + 2/*λ* and *I*_2_ = 2*λ* + 1/*λ*^2^ with *∂*_*λ*_*I*_1_ = 2 *λ* −2/*λ*^2^ and *∂*_*λ*_*I*_2_ = 2 −2/*λ*^3^. Using the zero normal stress condition, *P*_22_ = *P*_33_ = 0, we obtain the explicit expression for the uniaxial stress, *P*_11_ = 2 [*λ*− 1/*λ*^2^] *∂ψ*/*∂I*_1_ + 2 [1 −1/*λ*^3^] *∂ψ*/*∂I*_2_, which we can write explicitly in terms of the network weights ***w*** and ***w***^*∗*^,

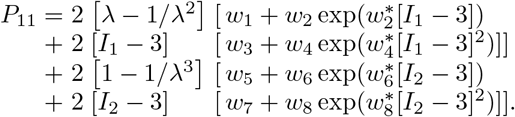

### Simple shear

In the shear experiment, we apply a torsion angle *ϕ*, that translates into the shear stress, *γ* = *r*/*Lϕ*, by multiplying it with the sample radius *r* and dividing by the initial sample length *L*. We can write the deformation gradient ***F*** in matrix representation as

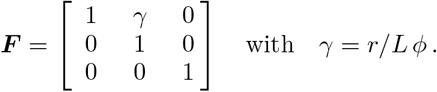

In shear, the first and second invariants and their derivatives are *I*_1_ = 3 + *γ*^2^ and *I*_2_ = 3 + *γ*^2^ with *∂*_*γ*_*I*_1_ = 2 *γ* and *∂*_*γ*_*I*_2_ = 2 *γ*. We obtain the explicit expression for the shear stress, *P*_12_ = 2 [*∂ψ*/*∂I*_1_ + *∂ψ*/*∂I*_2_] *γ*, which we can write explicitly in terms of the network weights ***w*** and ***w***^*∗*^,

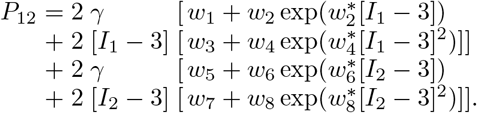

### Loss function

Our constitutive neural network learns the network weights, ***w*** and ***w***^*∗*^, by minimizing the loss function *L* that penalizes the mean squared error, the *L*_2_-norm of the difference between model and data, divided by the number of data points in tension, compression, and shear [28],

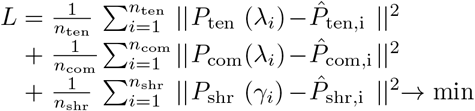

We train the network by minimizing the loss function with the ADAM optimizer, a robust adaptive algorithm for gradient-based first-order optimization using the tension, compression, and shear data for all eight meat products from Table 2.

### Best-in-class modeling

Instead of looking for the best possible fit of the models to the experimental data, we seek to discover meaningful constitutive models that are *interpretable* and *generalizable* [48], models that have a sparse parameter vector with a limited number of terms [49]. From combinatorics, we know that our network can discover 2^8^ −1 = 255 possible models, 8 with a single term, 28 with two, 56 with three, 70 with four, 56 with five, 28 with six, 8 with seven, and 1 with all eight terms. For practical purposes, we focus on the the 8 one-term models and the 28 two-term models, set all other weights to zero, and discover the non-zero weights that characterize the active terms. We summarize the results in a color-coded 8 ×8 error plot, as the average of the mean squared error across the tension, compression, and shear data, and report the parameters of the best-in-class one- and two-term models in Table 1. Motivated by the notable differences in the discovered models for tensile stretches up to 10% versus 35% in Figure 9, we decide to use the full tension data, not just the 10% stretch.

**Fig. 9.**
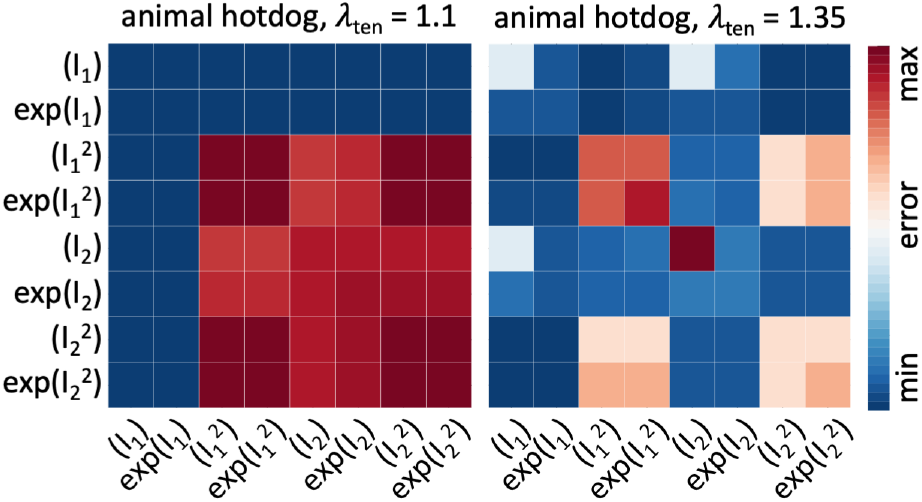
Model discovery for animal hotdog. Discovered one-term models, on the diagonal, and two-term models, off-diagonal, using tensile stretches up to 10% versus up to the peak stress of 35%. All models are made up of eight functional building blocks: linear, exponential linear, quadratic, and exponential quadratic terms of the first and second strain invariants *I*_1_ and *I*_2_. The color code indicates the quality of fit to the tension, compression, and shear data from Table 2, ranging from dark blue, best fit, to dark red, worst fit. The larger stretch range of 35% provides a clearer distinction of the quality of fit for the individual models.

### Food texture survey

We prepare bite-sized samples of the eight products, five plant-based, tofurky, plant-based sausage, plant-based hotdog, extrafirm tofu, and firm tofu, and three animal-based, spam turkey, animal sausage, and animal hotdog. We bake all samples in an oven until the plant-based products sre sufficiently warm, and the animal products reach a safe internal temperature for eating. We do not add any sauces or condiments, except for a small bit of oil to prevent excess sticking to the parchment paper. We keep the samples warm until serving. We recruit *n* = 16 participants to participate in three surveys: the ten-question Food Neophobia Survey [35] and the sixteen-question Meat Attachment Questionnaire [36], see Figure 10, and our own Food Texture Survey. We instruct each participant to eat a sample of each meat product and rank its texture features according to our survey. The survey uses a 5-point Likert scale with twelve questions. Each question starts with “this food is …”, followed by one of the following features [21, 22]: *soft, hard, brittle, chewy, gummy, viscous, springy, sticky, fibrous, fatty, moist*, and *meat-like*. The scale ranges from 5 for strongly agree to 1 for strongly disagree. This research was reviewed and approved by the Institutional Review Board at Stanford University under the protocol IRB-75418.

**Fig. 10.**
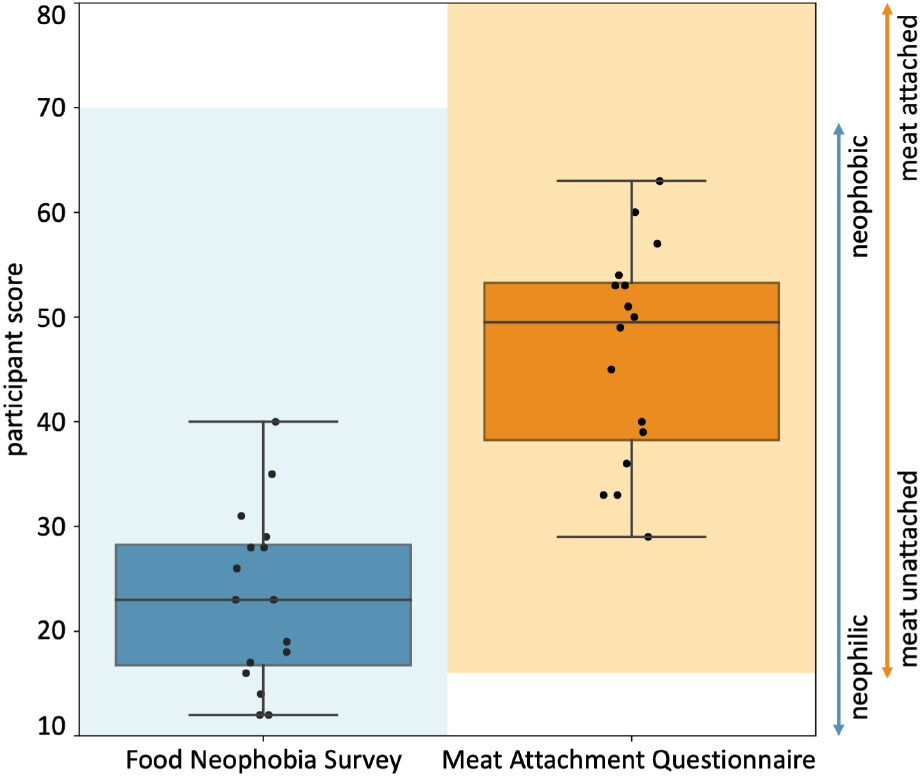
Participant scores for the Food Neophobia Survey and Meat Attachment Questionnaire. The Food Neophobia Survey uses a 7-point Likert scale with 10 questions so the light blue shaded range goes from 10 (neophilic, open to trying new foods) to 70 (neophobic, not open to trying new foods) [35]. The Meat Attachment Questionnaire uses a 5-point Likert scale with 16 questions so the light orange range goes from 16 (unattached to eating meat) to 80 (very attached to eating meat) [36]. The box-and-whisker plots of the minimum, first quartile, median, third quartile, and maximum participant scores are plotted in dark blue and dark orange for the Food Neophobia Survey and Meat Attachment Questionnaire, respectively. The black dots show individual scores.

## Data availability

The data are available in Table 2 as well as at https://github.com/LivingMatterLab/CANN.

## Code availability

The constitutive network code is available at https://github.com/LivingMatterLab/CANN.

## Acknowledgments

This project was supported by the NSF Graduate Research Fellowship to SS, by the Emmy Noether Grant 533187597 to KL, and by the NSF CMMI Award 2320933 and the ERC Advanced Grant 101141626 to EK.

## Authors’ contributions

SS designed the 3D printed grips, contributed to the code, conducted the analyses, and wrote the paper. SS and ED designed and oversaw the testing protocols. DA, MA, AD, RD, YL, MPV, and VPM performed the experimental testing. KL wrote the constitutive neural network code. ML and EK conceptualized the project and edited the paper.

## Competing interests

The authors declare no competing interests.

## Notes

### Competing Interest Statement

The authors have declared no competing interest.

### Summary of Updates

slight adjustment in order of methods/results and change of two words in the title

